# Full-assembly screening reveals mobile antibiotic-resistance cargo missed by chromosome-only genomes

**DOI:** 10.64898/2026.06.11.731541

**Authors:** Saniya, Abdullah Ahmad Khan

**Author notes:** Corresponding author: Abdullah Ahmad Khan.

## Abstract

**Background:** Probiotic bacteria occupy the same gut niches as enteric pathogens, prompting concern that probiotic strains might carry or contribute mobile antibiotic-resistance genes (ARGs). Genome-based screening is routinely used to assess this risk, but many screens use chromosome-level assemblies that may omit plasmid-borne, high-mobility cargo. We quantified this effect and compared the mobile context resistomes of probiotic-associated and pathogen reference genomes.

**Methods:** We screened 50 bacterial reference genomes (25 probiotic-associated, 25 pathogen/comparator) using a reproducible workflow with the CARD nucleotide catalog, PlasmidFinder replicons, and ISfinder insertion sequences. Each ARG was assigned a fourtier *in silico* mobile-context risk category from plasmid co-localization and insertion-sequence (IS) flanking. The identical strain panel was screened in matched full-assembly and chromosome-only modes. Acquired calls were curated against intrinsic/efflux/biocide determinants and cross-checked with AMRFinderPlus, ResFinder, and targeted BLAST, with MOB-suite as a plasmid/mobility overlay.

**Results:** Full-assembly screening detected 373 ARG loci versus 338 in chromosome-only mode on the same strains, increasing High-risk calls from 4 to 15 and recovering 32 plasmid replicons (chromosome-only: 0). All 15 High-risk mobile-context loci occurred in pathogen/comparator genomes and none in probiotic-associated genomes; no ARG was shared across groups at ≥95% nucleotide identity (0/175 edges). Per-strain ARG burden was higher in pathogen genomes (mean 12.32 versus 0.52 loci; Mann-Whitney *U* = 606.5, *P*< 0.001). Most priority High-risk loci were corroborated by one or more external tools, with discordant calls retained explicitly as flagged records.

**Conclusions:** Chromosome-only screening materially undercounts mobile ARG cargo. In this reference-genome panel, high-risk mobile-context loci were concentrated in pathogen/comparator genomes — an *in silico* reference-genome-level safety signal rather than evidence about commercial products or genetic transfer.

**Data Summary:** No new sequencing data were generated; all genomes are publicly available reference assemblies from NCBI RefSeq.

1. Genome accessions for all 50 genomes are in Supplementary Table S1.
2. Source code (screening, validation, and figure scripts, including `make_figures.py`) is available at https://github.com/abdullahak07/1dprob.
3. Processed result tables are summarised in Supplementary Tables S2–S10.
4. External validation outputs (AMRFinderPlus, ResFinder, BLAST, MOB-suite) are provided as supplementary data.
5. Tool/database versions and detection thresholds are in Supplementary Table S9. The authors confirm that all supporting data, code, and protocols are available within the article, the cited repositories, or the supplementary material.

**Impact Statement:** Genome-based screening is widely used to judge whether a bacterial strain carries transferable antibiotic-resistance genes, including in the safety assessment of probiotic-associated species. We show that the assembly level chosen for such screening materially affects the result: on an identical panel of 50 reference genomes, chromosome-only analysis recovered fewer than a third of the high-risk, mobile-context resistance loci detected when plasmid replicons were included. Applying full-assembly screening with transparent, multi-tool external validation, high-risk mobile-context resistance genes were concentrated in pathogen/comparator genomes and absent from the probiotic-associated genomes in this panel, with no cross-group sharing at high identity. These observations argue for full-assembly inputs and explicit mobile-context interpretation in genome-based resistance screening and provide a cautious, reference-genome-level safety signal; they are not claims about commercial products and do not demonstrate genetic transfer.

## Introduction

Antimicrobial resistance (AMR) spreads not only by clonal expansion but by horizontal movement of resistance genes on plasmids and other mobile genetic elements (MGEs). The clinical risk posed by an ARG therefore depends heavily on its genomic context: a determinant on a plasmid or flanked by insertion sequences (IS) has greater mobilization potential than the same gene fixed in the core chromosome. This context-dependence is central to interpreting AMR in any genome, and especially in organisms deliberately introduced into the human gut.

Probiotic-associated species—lactobacilli, bifidobacteria, enterococci, and related taxa— are consumed in large numbers and share the gut environment with enteric pathogens, motivating interest in whether they carry, or could disseminate, transferable resistance genes. Genome-based screening with curated catalogs, such as CARD, and tools, such as AMRFinderPlus and ResFinder, is now routine. However, screening outcomes depend on inputs that are not always reported. Many analyses use chromosome-level or draft assemblies in which plasmid replicons are incomplete or absent; because mobile resistance cargo is enriched on plasmids, such analyses may systematically under-represent the high-mobility determinants of greatest interest. Recent plasmid-resolved and mobilome-focused studies in *Microbial Genomics* similarly show that plasmid-aware analysis can materially affect interpretation of resistance carriage, surveillance, and genome-based safety assessment.

We examine this effect directly. Using a balanced panel of 50 reference genomes (25 probiotic-associated, 25 pathogen/comparator), we screened for ARGs and their plasmid and IS contexts, assigned a four-tier *in silico* mobile-context risk category, and ran the same strains in matched full-assembly and chromosome-only modes. We curated acquired calls separately from intrinsic, efflux and biocide determinants, cross-validated identities with AMRFinderPlus, ResFinder and targeted BLAST, and overlaid plasmid/MGE mobility from MOB-suite. Our central question is methodological: how much does the choice of assembly inputs change the picture of mobile-context resistance? The comparative biology follows as an application.

## Methods

### Genome selection

Fifty bacterial reference genomes were analyzed: 25 probiotic-associated (well-characterized type or reference strains of species used in probiotic and fermented-food contexts) and 25 pathogen/comparator genomes (clinically relevant species and reference isolates), obtained from NCBI RefSeq. Where available, full assemblies comprising the chromosome plus all RefSeq plasmid replicons were retrieved; otherwise, the primary chromosomal replicon was used. All genomes are reference assemblies, not isolates from commercial probiotic products. Accessions and metadata are in Supplementary Table S1.

### ARG detection

Acquired ARGs were detected against the CARD nucleotide catalogue using a *k*-mer-indexed ungapped alignment, retaining loci at ≥95% nucleotide identity and ≥80% reference coverage. Overlapping allele models were collapsed to one best locus per reciprocal-overlap cluster (overlap ≥0.8; best by identity, then coverage, then length) to avoid inflating counts. Allele names are reported as best database matches and treated as confirmed only where supported by external tools.

### Mobile-genetic-element detection

Plasmid replicons were detected against PlasmidFinder and insertion sequences against ISfinder using the same engine at ≥80% identity and ≥60% coverage. Conjugation-system markers were screened by nucleotide similarity; none were detected, and conjugative status was not inferred from the in-house screen.

### Mobile-context risk classification

Each ARG was assigned a four-tier category based on genomic context. High: plasmid-associated and/or IS-flanked on both sides within 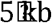. Medium: weaker mobile-context support. Low: acquired ARGs without high-risk context. Negligible: intrinsic determinants. This is an *in silico* mobile-context heuristic for triage, not an experimentally calibrated predictor of transfer, and is interpreted accordingly.

### Curation

Raw ARG calls were preserved. For interpretation, calls were classified as acquired mobile ARG, intrinsic/core gene, efflux/regulatory marker, biocide/stress marker, or ambiguous using a documented rule set, so that intrinsic, efflux, and biocide determinants were not over-interpreted as acquired cargo. Only acquired and ambiguous categories were included in the mobile-ARG analysis; raw calls are in Supplementary Tables S2–S3.

### External identity validation

Identities were cross-checked with AMRFinderPlus, ResFinder and targeted BLAST. A locus was counted as externally confirmed only when an external hit matched on strain and contig, overlapped the in-house coordinates, and named a compatible gene family under a normalization reconciling catalog synonyms (e.g. *bla*_CTX-M-14_/CTX-M-14, *erm*(B)/ErmB, *strA*/*aph(3)-Ib*), at ≥90% identity and ≥80% coverage where available. Loci were assigned confirmed by a single named tool, confirmed by ≥2 tools, discordant (an overlapping external hit naming a conflicting family), or not checked. Discordant calls were retained and flagged for review (Supplementary Table S10).

### Mobility overlay

Plasmid/MGE mobility context for High/Medium loci was overlaid with MOB-suite to corroborate plasmid contig assignment and mobilization potential. MOB-suite was the only external mobility tool used. A second tool, geNomad, was evaluated but did not complete within the available compute environment and is not part of the reported result; no geNomad-derived claim is made.

### Statistics

Per-strain mobile ARG burden was compared with the Mann-Whitney *U* test. Group differences in the proportion of loci that were High/Medium, plasmid-associated, or IS-flanked on both sides were tested with Fisher’s exact test at the locus level. Tests were two-sided; *P*-values are interpreted cautiously given the small number of probiotic ARG loci.

### Implementation and reproducibility

The core screening workflow was implemented in Python with lightweight dependencies, while external validation was performed using AMRFinderPlus, ResFinder, BLAST, and MOB-suite. Source code, result tables, validation files, figure scripts, the accession table, and the version/threshold record are deposited (Data Summary; Supplementary Table S9).

## Results

### Dataset overview

Across the 50 reference genomes, full-assembly screening detected 373 ARG loci (15 High-risk, 43 Medium-risk, 315 Low-risk), 32 plasmid replicons, 1116 IS features, and 0 conjugation markers (Table 1; workflow in Fig. 1). Gene-sharing analysis at ≥95% nucleotide identity yielded 175 edges, none connecting a probiotic-associated to a pathogen/comparator genome.

Summary of the reference-genome panel and aggregate outputs.

| Item | Value |
| --- | --- |
| Probiotic-associated reference genomes | 25 |
| Pathogen/comparator reference genomes | 25 |
| Total ARG loci | 373 |
| High-risk / Medium-risk / Low-risk | 15 / 43 / 315 |
| Plasmid replicons | 32 |
| Insertion-sequence features | 1116 |
| Conjugation markers | 0 |
| Gene-sharing edges ($\geq 95\%$ identity) | 175 |
| Probiotic–pathogen (cross-group) edges | 0 |

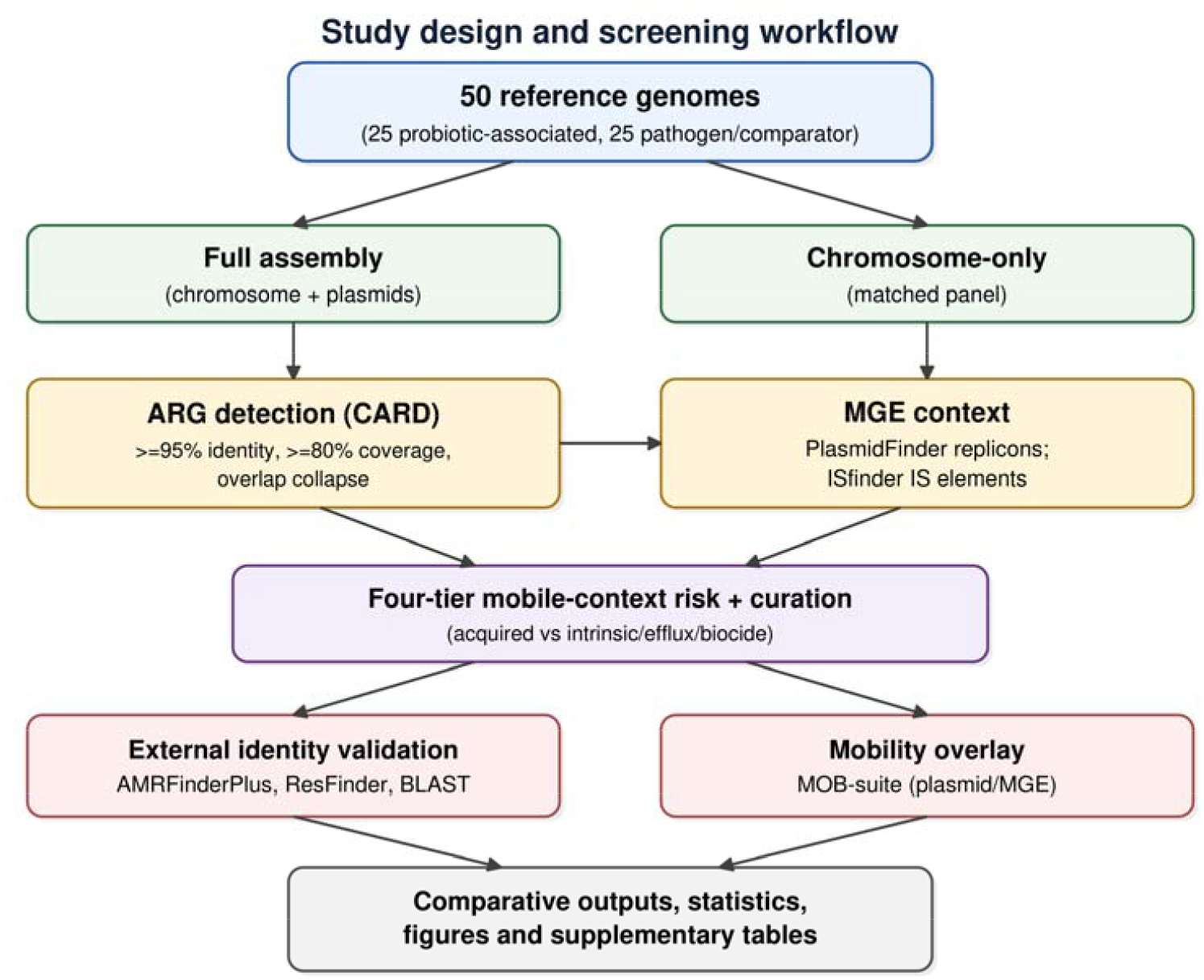
Study design and screening workflow. Fifty reference genomes (25 probiotic-associated, 25 pathogen/comparator) were screened in matched full-assembly and chromosome-only modes for ARGs (CARD) and MGE context (PlasmidFinder replicons; ISfinder IS), assigned a four-tier mobile-context risk category, curated (acquired versus intrinsic/efflux/biocide), and externally validated for identity (AMRFinderPlus, ResFinder, BLAST) and mobility (MOB-suite).

### Full-assembly versus chromosome-only screening

On the identical panel, chromosome-only screening detected 338 ARG loci (4 High-risk, 21 Medium-risk, 313 Low-risk) and 0 plasmid replicons, with 1003 IS features. Full-assembly screening recovered a net 35 additional ARG loci and changed the mobile-context picture: High-risk calls increased from 4 to 15, Medium-risk from 21 to 43, plasmid replicons from 0 to 32, and IS features from 1003 to 1116 (Table 2; Fig. 2). The additional High- and Medium-risk loci were predominantly plasmid-associated. The net change in loci is a count of detected ARG rows and equals the sum of the per-tier increases (Low +2, Medium +22, High +11); a gene recovered at more than one plasmid locus in a strain contributes additional rows without representing additional gene–strain combinations.

Matched full-assembly versus chromosome-only comparison on the identical 50-strain panel.

| Metric | Chromosome-only | Full assembly | Delta |
| --- | --- | --- | --- |
| ARG loci | 338 | 373 | +35 |
| High-risk | 4 | 15 | +11 |
| Medium-risk | 21 | 43 | +22 |
| Low-risk | 313 | 315 | +2 |
| Plasmid replicons | 0 | 32 | +32 |
| Insertion-sequence features | 1003 | 1116 | +113 |

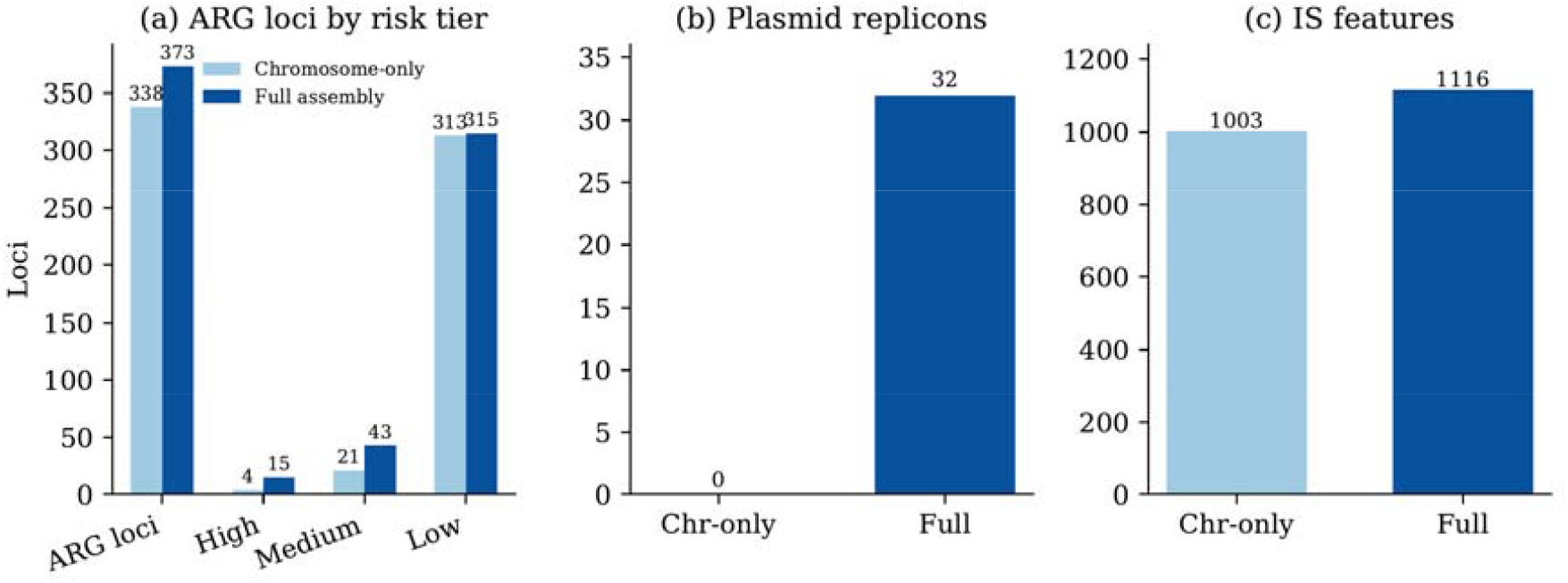
Matched full-assembly versus chromosome-only comparison. (a) ARG loci by risk tier; (b) plasmid replicons; (c) insertion-sequence features, for the identical 50-strain panel. High-risk loci increase from 4 to 15 and plasmid replicons from 0 to 32 under full assembly.

### ARG and risk distribution by group

The strain-resolved heatmap showed a clear separation in ARG burden and mobile-context risk between probiotic-associated and pathogen/comparator genomes (Fig. 3). All 15 High-risk mobile-context loci were detected in pathogen/comparator genomes, but none were detected in the probiotic-associated group. Per-strain mobile ARG burden was higher in pathogen/comparator genomes than in probiotic-associated genomes (mean 12.32 versus 0.52 loci; Mann-Whitney *U* = 606.5, *P* < 0.001). Many probiotic-associated reference genomes carried no acquired mobile ARGs, supporting the interpretation that high-risk mobile-context resistance cargo was concentrated in the pathogen/comparator group in this reference panel.

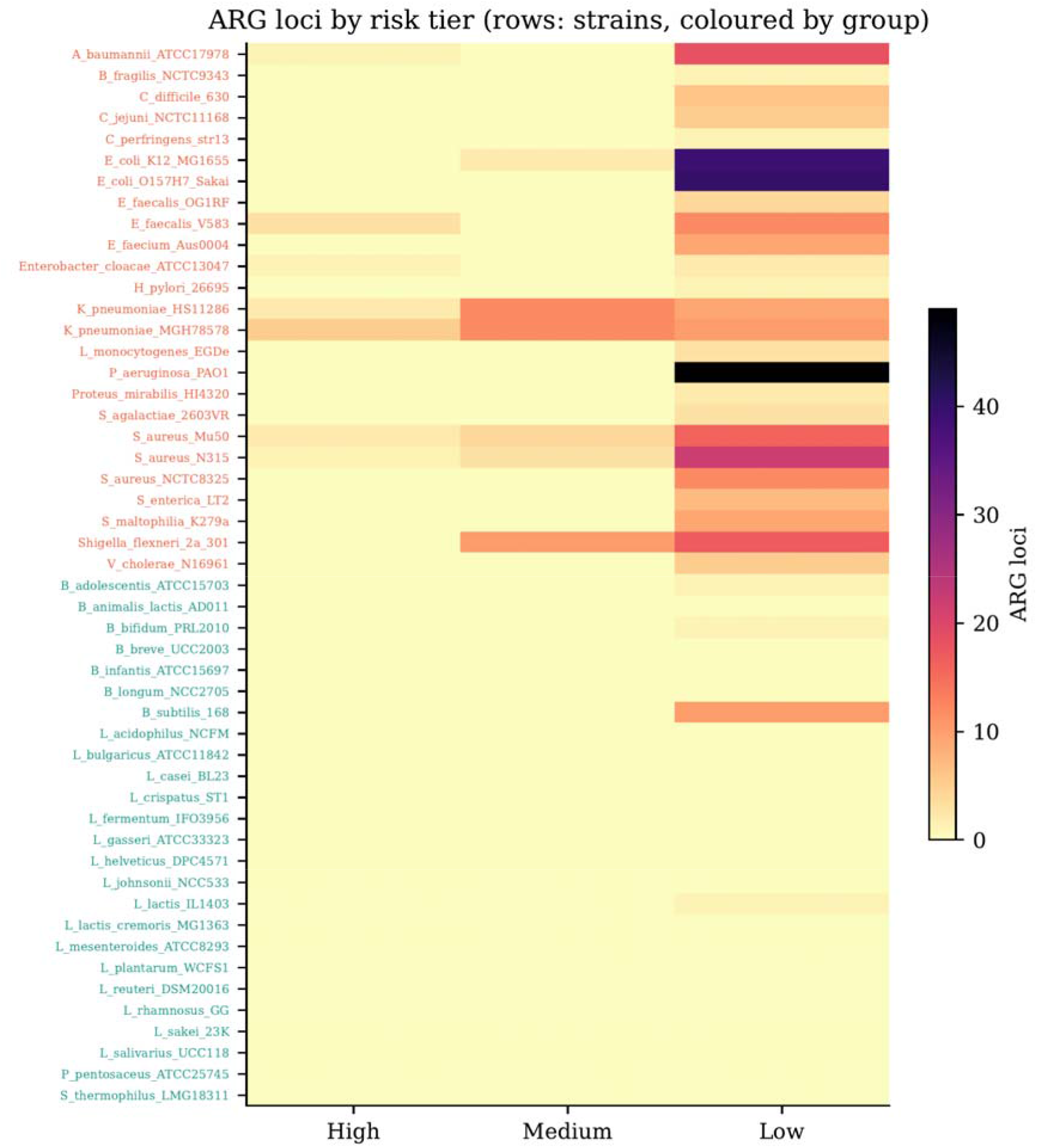
Strain-resolved ARG burden and mobile-context risk by group. Heatmap showing ARG burden and risk-tier distribution across individual reference genomes. Rows represent genomes grouped as probiotic-associated or pathogen/comparator, and columns represent ARG/risk features. High mobile-context loci were detected only in pathogen/comparator genomes, whereas probiotic-associated reference genomes carried few ARG loci and no High-risk calls.

### Curation

Of the 373 loci, curation classified 316 as acquired mobile ARG, 39 as efflux/regulatory, 9 as intrinsic/core, 5 as ambiguous, and 4 as biocide/stress (Supplementary Fig. S1). Separating intrinsic, efflux, and biocide determinants reduced the High/Medium acquired-cargo set from 58 raw High/Medium loci to 45 curated acquired-cargo High/Medium loci. Raw calls were retained throughout (Supplementary Tables S2–S4).

### External identity validation

ARG identity validation imported AMRFinderPlus, ResFinder, and targeted BLAST, confirming 94, 68, and 46 loci respectively; 68 loci were confirmed by two or more external identity tools, while 33 discordant loci were retained as flagged calls for review, and 224 loci were not externally checked or were retained as best database matches (Table 3; Supplementary Fig. S2). We do not claim that all 373 loci were externally confirmed. Discordant calls may reflect nomenclature, family-boundary or database-version differences between tools, but were retained as flagged calls pending manual review (Supplementary Table S10) rather than silently reconciled.

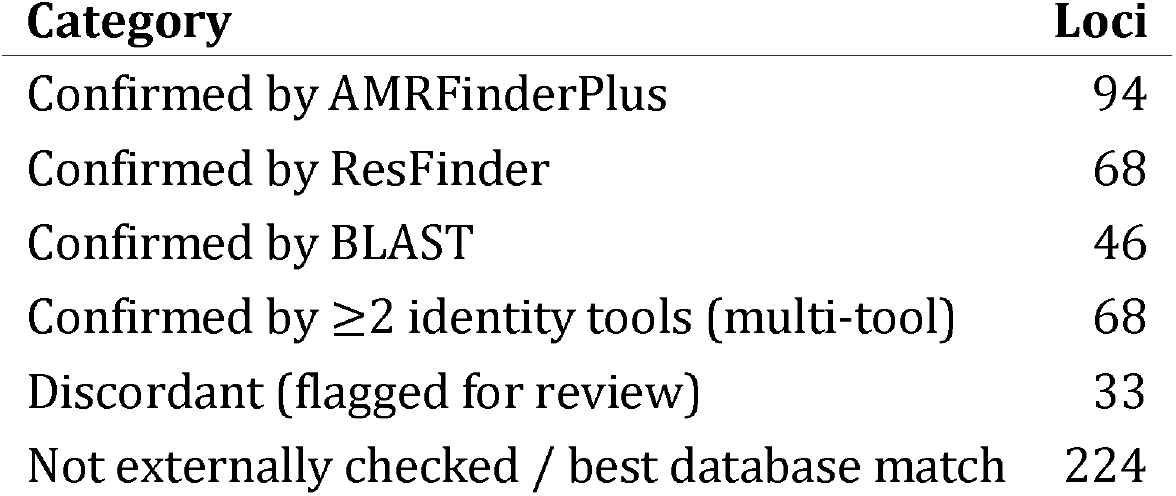
External validation summary. Counts are loci; a locus may be confirmed by more than one tool.

| Category | Loci |
| --- | --- |
| Confirmed by AMRFinderPlus | 94 |
| Confirmed by ResFinder | 68 |
| Confirmed by BLAST | 46 |
| Confirmed by $\geq 2$ identity tools (multi-tool) | 68 |
| Discordant (flagged for review) | 33 |
| Not externally checked / best database match | 224 |

Most High-risk loci were corroborated by multiple tools; exceptions were retained with explicit validation status. For example, *erm*(B) in *Enterococcus faecalis* V583, bla_CTX-M-14_ in *Klebsiella pneumoniae* HS11286, *sul2* in *K. pneumoniae* MGH78578 and *sul2* in *Acinetobacter baumannii* ATCC 17978 were each confirmed by AMRFinderPlus, ResFinder and BLAST (Table 4; Fig. 4).

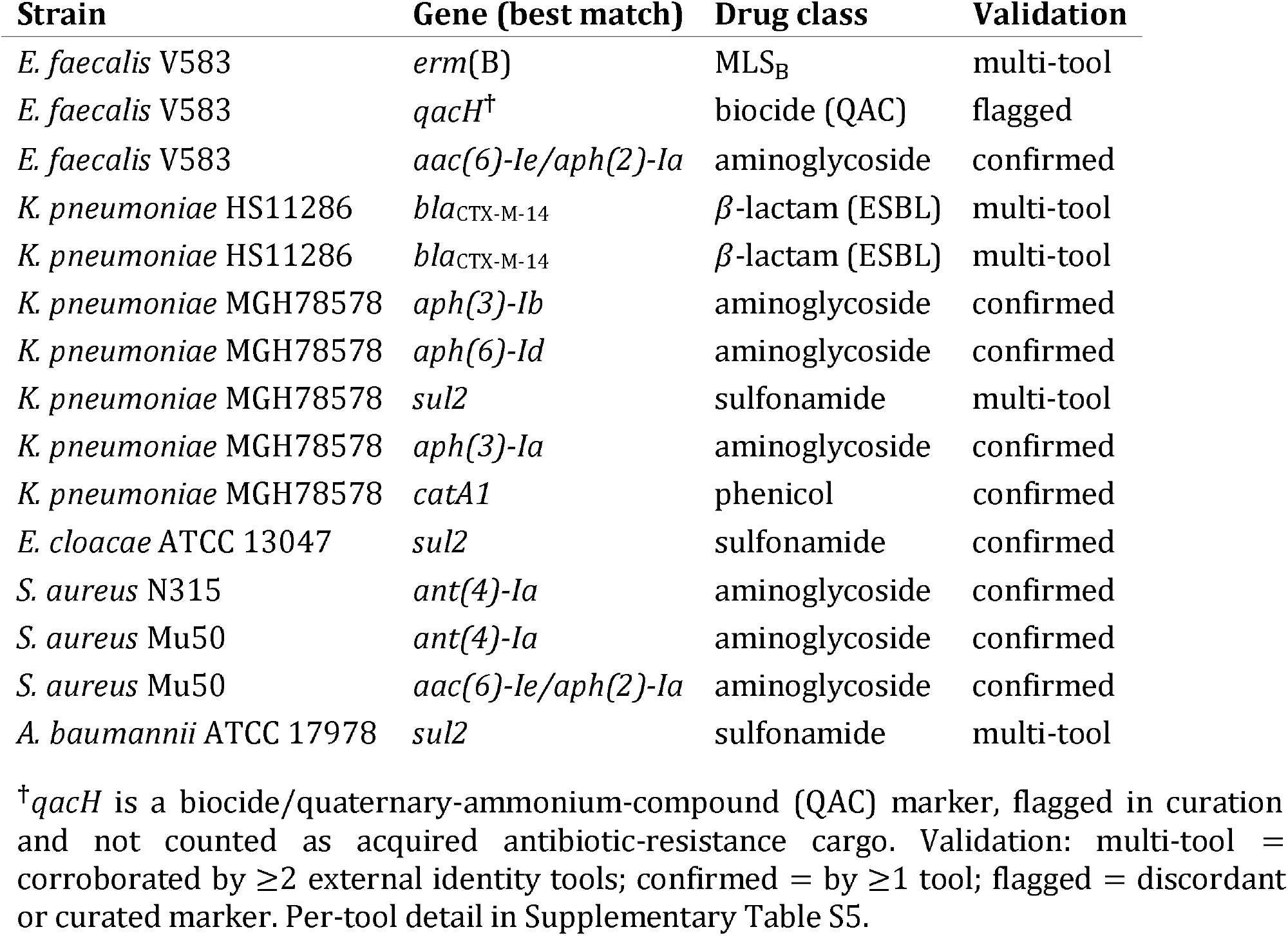
High-risk mobile-context loci, detected only in pathogen/comparator genomes. All satisfied the High criterion (IS flanking on both sides; a subset additionally plasmid-associated, e.g. bla_CTX-M-14_). Contigs, coordinates and per-tool validation are in Supplementary Tables S4– S5.

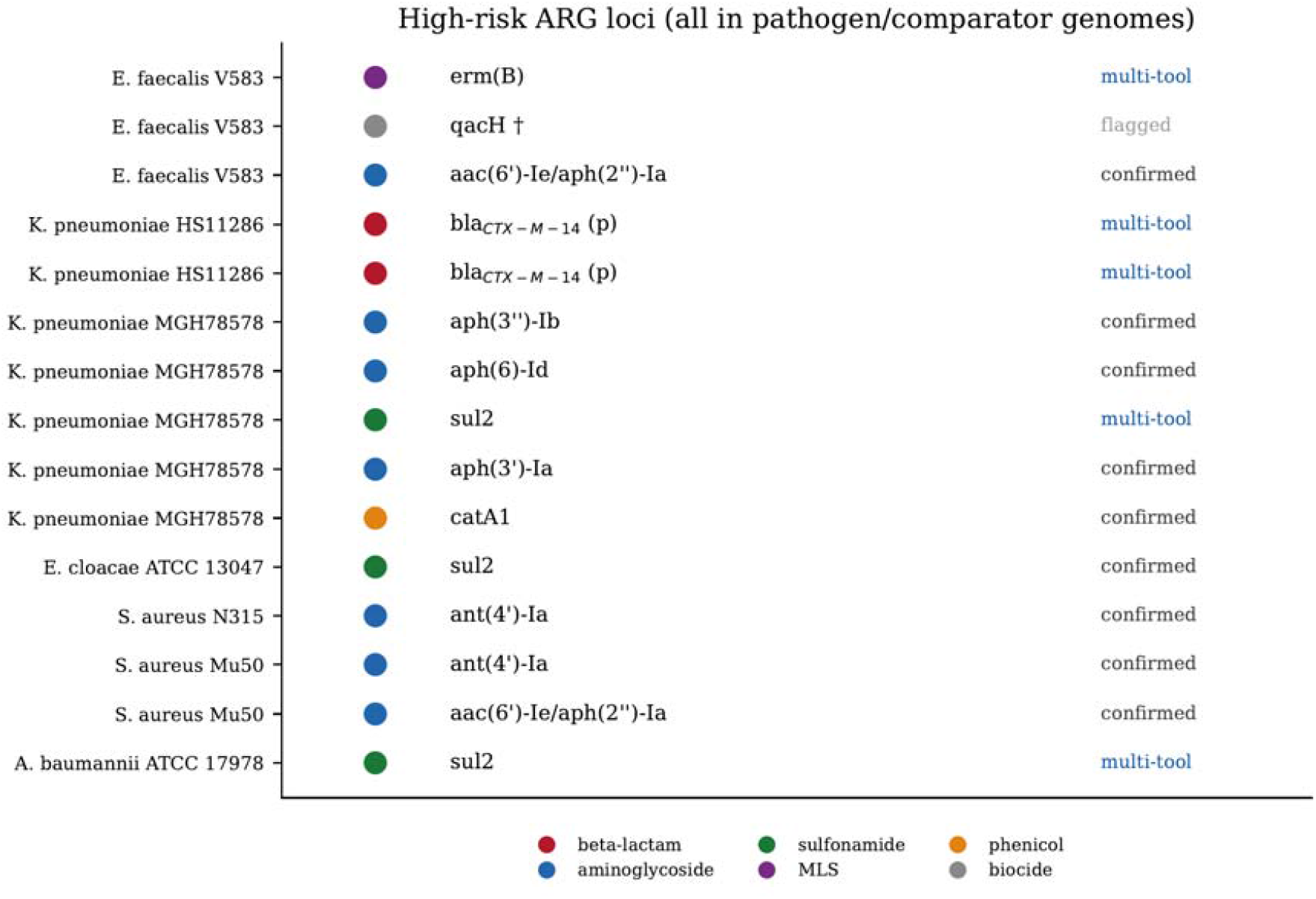
High-risk ARG loci and validation status. The 15 High-risk mobile-context loci, all in pathogen/comparator genomes, coloured by drug class and labelled with external validation status (multi-tool / confirmed / flagged). “(p)” marks plasmid-associated loci; qacH (†) is a biocide marker flagged in curation.

### Mobility overlay

Of the 58 High/Medium loci, 33 were on plasmid contigs, 15 were IS-flanked on both sides, and 37 were positive for *in silico* mobile-context potential; 33 were corroborated as plasmid/MGE-associated by the MOB-suite overlay (Supplementary Fig. S3). No locus was externally predicted to be conjugative, consistent with the absence of conjugation markers. Mobility results are reported as mobile-context potential, not demonstrated mobilization.

### High-risk loci

High-risk loci were detected only in pathogen/comparator genomes (Table 4; Fig. 4), including *erm*(B), *qacH* and *aac(6)-Ie/aph(2)-Ia in E. faecalis V583; bla*_CTX-M-14_ on plasmid-associated loci in *K. pneumoniae* HS11286; *aph(3)-Ib, aph(6)-Id, sul2, aph(3)-Ia* and catA1 in K. pneumoniae MGH78578; *sul2 in Enterobacter cloacae* ATCC 13047; *ant(4)*-Ia and *aac(6)*-*Ie/aph(2)-Ia* in *Staphylococcus aureus* N315/Mu50; and *sul2* in A. *baumannii* ATCC 17978.

### Gene sharing

The gene-sharing network comprised 175 edges at ≥95% nucleotide identity, all within the pathogen/comparator group; no edge connected a probiotic-associated to a pathogen/comparator genome (Supplementary Fig. S5).

### Statistics

Per-strain mobile ARG burden was higher in pathogen/comparator than in probiotic-associated genomes (Mann– Whitney *U* = 606.5, P< 0.001; means 12.32 versus 0.52 loci). Locus-level enrichment tests were not formally significant: High/Medium risk (Fisher’s exact *P*= 0.38), plasmid-associated (*P* = 0.61), and IS-flanked-both-sides (*P* = 1). The per-strain burden difference is therefore substantial and in the expected direction, whereas locus-level enrichment is underpowered because probiotic-associated genomes contributed very few ARG loci; we describe the high-risk pattern as an observed concentration in pathogen/comparator genomes rather than as statistically significant enrichment.

## Discussion

The principal result is methodological. On a fixed 50-genome panel, the assembly level used for screening substantially changed the picture of mobile-context resistance: chromosome-only analysis recovered four High-risk loci and no plasmid replicons, whereas full-assembly analysis of the same strains recovered fifteen High-risk loci and thirty-two replicons. Because high-mobility resistance cargo is enriched on plasmids and IS-associated regions, chromosome-only or plasmid-incomplete inputs can omit precisely the determinants that screening is meant to flag, consistent with recent plasmid-resolved and mobilome-focused genomic studies. This has practical implications for any genome-based resistance assessment, including the safety screening of probiotic-associated species, and argues for full-assembly inputs and explicit mobile-context interpretation wherever feasible.

Applying this workflow, high-risk mobile-context ARGs were absent from the probiotic-associated reference genomes in this panel and were concentrated in the pathogen/comparator genomes, with no ARG shared between groups at ≥95% nucleotide identity. We interpret this as a cautious, *in silico*, reference-genome-level safety signal for the probiotic-associated set analyzed here. We deliberately do not extend it further: these are reference genomes, not isolates from commercial products, and an absence of shared high-identity ARGs *in silico* is not evidence about genetic transfer, which we did not assay.

External validation supported the priority calls while exposing their limits. Many loci, including the principal High-risk determinants, were corroborated by two or more independent tools; a minority were discordant. We retained discordant calls as flagged entries rather than discarding them. Discordant calls may reflect nomenclature, family-boundary or database-version differences between tools, but were retained as flagged calls pending manual review rather than reconciled by assumption. Reporting confirmed, multi-tool-confirmed, discordant and unconfirmed categories separately is, in our view, more useful than a single headline “validated” count.

We emphasize that the four-tier risk scheme is a triage heuristic based on genomic context, not a calibrated predictor of transfer. A High-risk classification denotes mobile-context potential, not demonstrated mobilization. Likewise, the per-strain burden difference, although strong, is in the expected direction and is not the novel element of this work; the contributions are the matched full-assembly versus chromosome-only quantification, the cautious mobile-context classification, and the transparent multi-tool validation and discordance handling.

### Scope and interpretation

First, the study uses public genomes and standard tools: the contribution is the matched quantification of how assembly input changes mobile-context results, with explicit validation and discordance reporting that are often omitted. Second, the risk heuristic is unvalidated: it is presented as a triage scheme, not a transfer predictor. Third, discordant calls might be seen as weakening the picture; we address this by flagging them transparently and retaining them for review. Fourth, that no product-level conclusion is possible: we make none. Fifth, a second mobility tool was not completed: MOB-suite served as the overlay, and no geNomad-based claim is made. Future work should extend the panel beyond 50 genomes, incorporate product-level isolate sequencing and metagenomics, and pair high-risk loci with experimental mobilization assays.

## Limitations

The genomes are reference assemblies, not isolates from commercial products, so the findings do not speak to product-level safety. All inferences are *in silico*; no conjugation or transfer experiments were performed, and the risk scheme is not experimentally calibrated. No new sequencing was generated. The panel of 50 genomes was selected for balance and characterization rather than as a global survey, limiting generalisability. Results depend on the versions of CARD, PlasmidFinder, ISfinder, AMRFinderPlus, ResFinder, and MOB-suite and on the chosen thresholds (Supplementary Table S9). A subset of calls remained discordant and are flagged rather than resolved, and 224 lower-priority loci were retained as best database matches without external confirmation. geNomad was not completed and is excluded. Conjugation markers were not detected by nucleotide screening, a relatively insensitive approach that should not be read as definitive absence.

## Author contributions

Using the CRediT taxonomy: **Saniya** — Conceptualization, Validation, Data curation. Investigation, writing original draft. **Abdullah Ahmad Khan** — Conceptualization, Methodology, Software, Validation, Formal analysis, Investigation, Writing review & editing, Visualization.

## Acknowledgements

The authors thank Murdoch University IT department for computing support. This study used publicly available reference genomes from NCBI RefSeq and the CARD, PlasmidFinder, and ISfinder resources, as well as the AMRFinderPlus, ResFinder, and MOB-suite tools; we acknowledge their developers and curators.

## Supplementary material

*Supplementary figures*. **S1** Curation categories. **S2** External identity-validation status. **S3** Mobility-context status for High/Medium loci. **S4** Per-strain ARG burden by group. **S5** Gene-sharing network coloured by group. S6 Per-strain ARG burden boxplot. S7 Drug-class distribution by group.

*Supplementary tables*. **S1** Strain metadata and accessions. **S2** Full ARG locus table (raw calls preserved). **S3** Curated ARG calls. **S4** Curated High/Medium acquired-cargo loci. **S5** External identity validation per locus. **S6** Mobility validation per locus. **S7** Gene-sharing edges. **S8** Statistical test outputs. **S9** Tool/database versions and detection thresholds. **S10** Discordant loci flagged for review.

**Supplementary Table S9:**
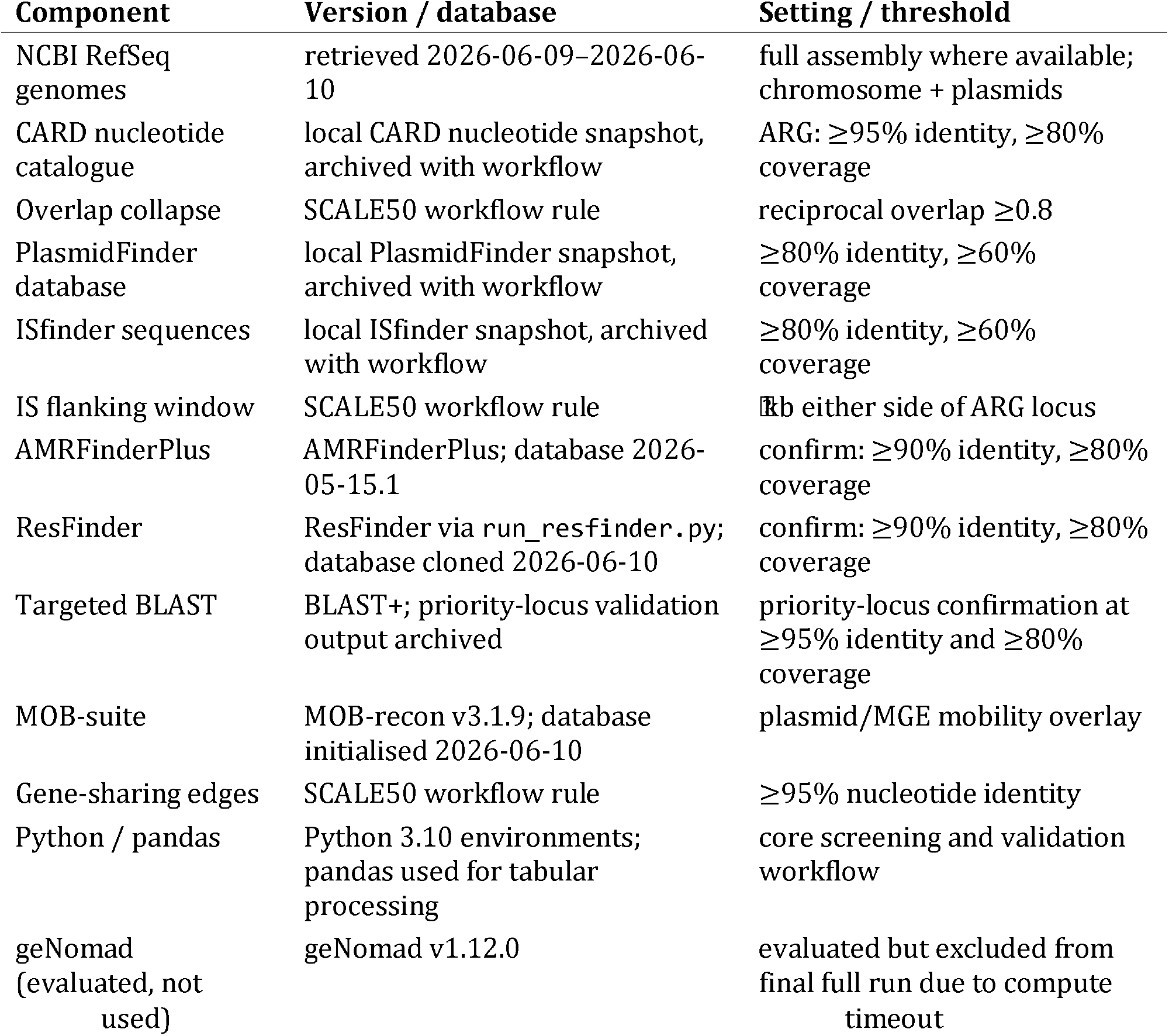
tool/database versions and detection thresholds used in the SCALE50 analysis. Database snapshots and external validation outputs should be archived with the supplementary data to ensure reproducibility.

## Supplementary figures

**Supplementary Fig. 1.**
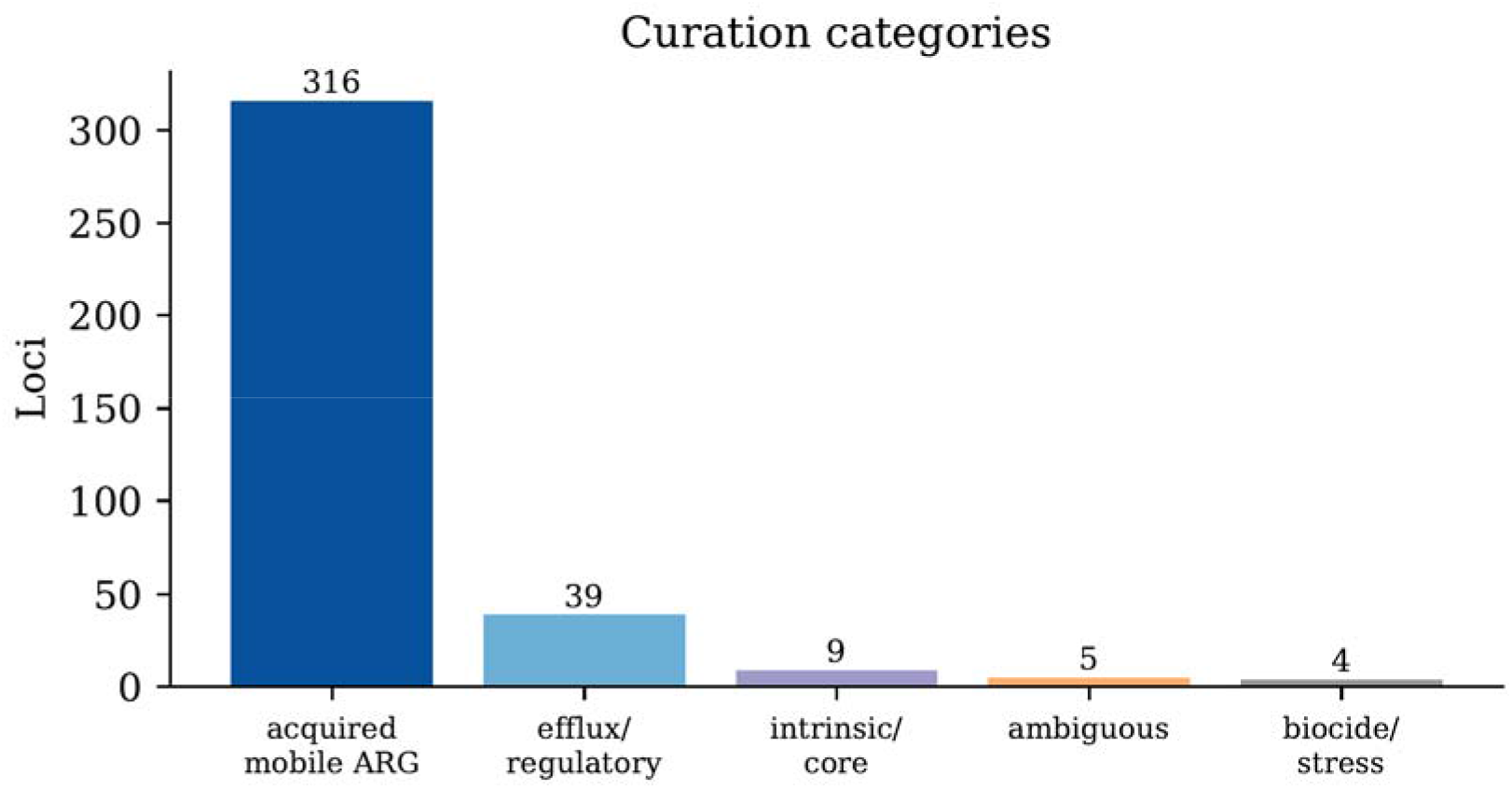
Curation categories. Distribution of the 373 ARG loci across curation categories: acquired mobile ARG (316), efflux/regulatory (39), intrinsic/core (9), ambiguous (5) and biocide/stress (4).

**Supplementary Fig. 2.**
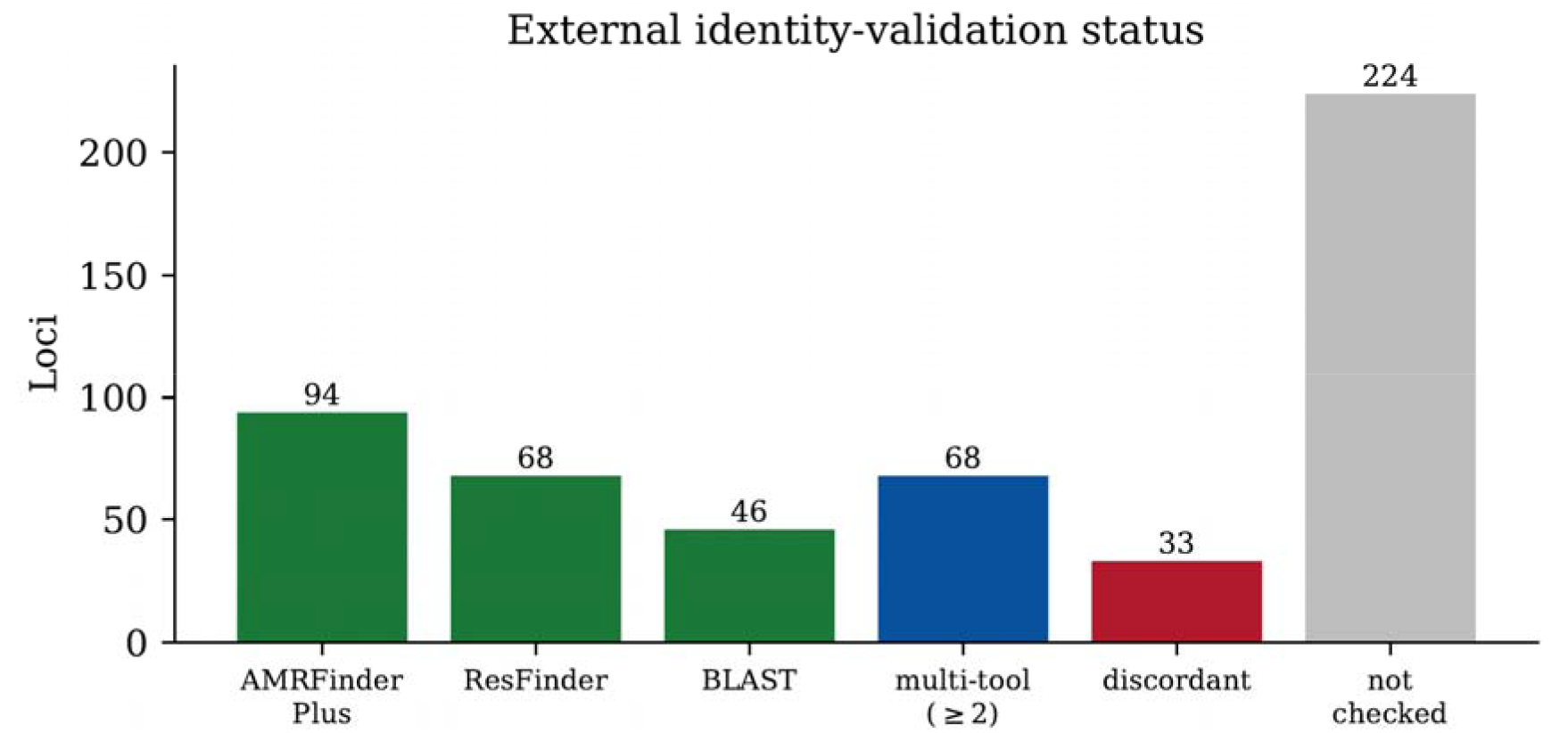
External identity-validation status. Per-tool confirmation counts (AMRFinderPlus, ResFinder, BLAST), multi-tool confirmation (≥2 tools), discordant calls retained for review, and loci not externally checked.

**Supplementary Fig. 3.**
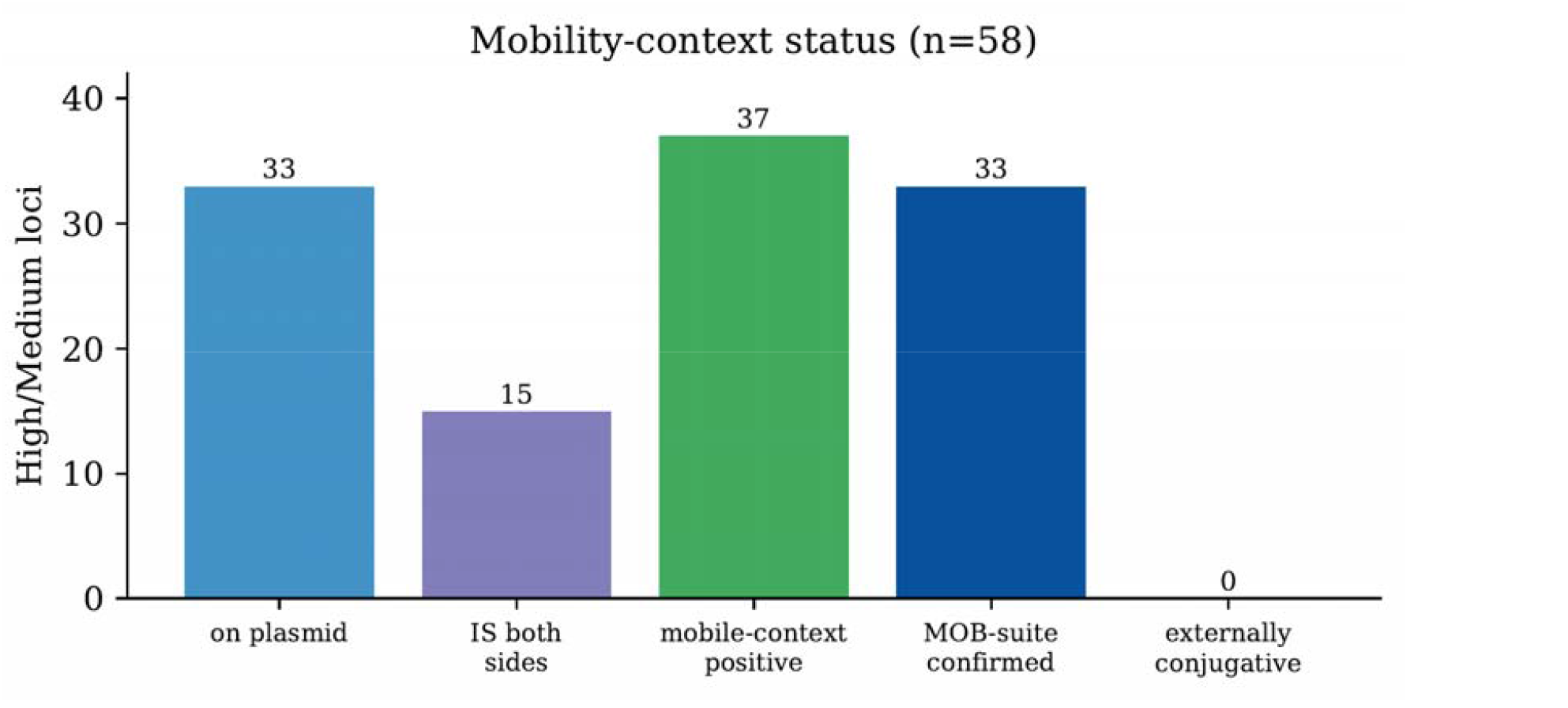
Mobility-context status. For the 58 High/Medium loci: plasmid-associated, IS-flanked on both sides, mobile-context positive, MOB-suite-confirmed, and externally predicted conjugative (none).

**Supplementary Fig. 4.**
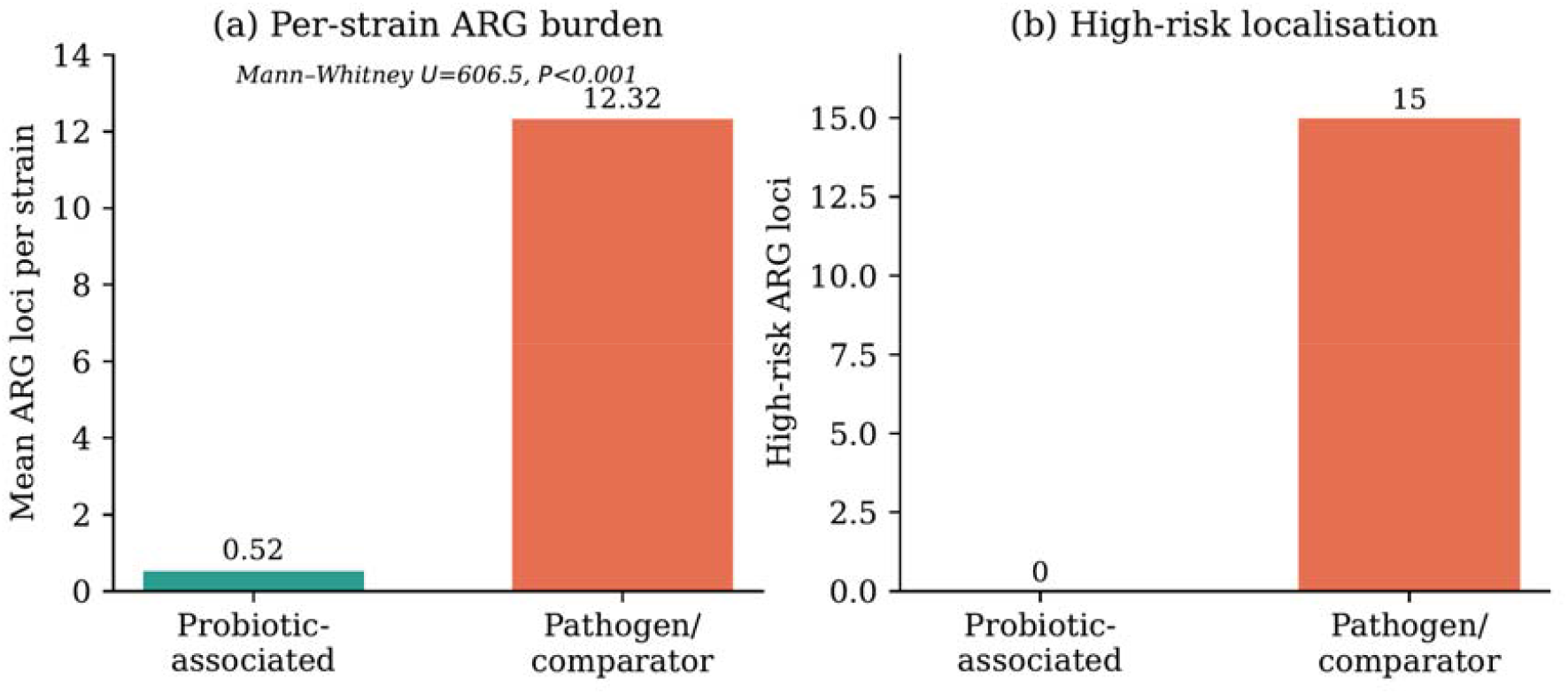
Per-strain ARG burden by group. Per-strain ARG burden in probiotic-associated and pathogen/comparator reference genomes. Pathogen/comparator genomes carried a higher ARG burden than probiotic-associated genomes (Mann–Whitney U = 606.5, P < 0.001), consistent with the concentration of mobile-context resistance cargo in the pathogen/comparator group.

**Supplementary Fig. 5.**
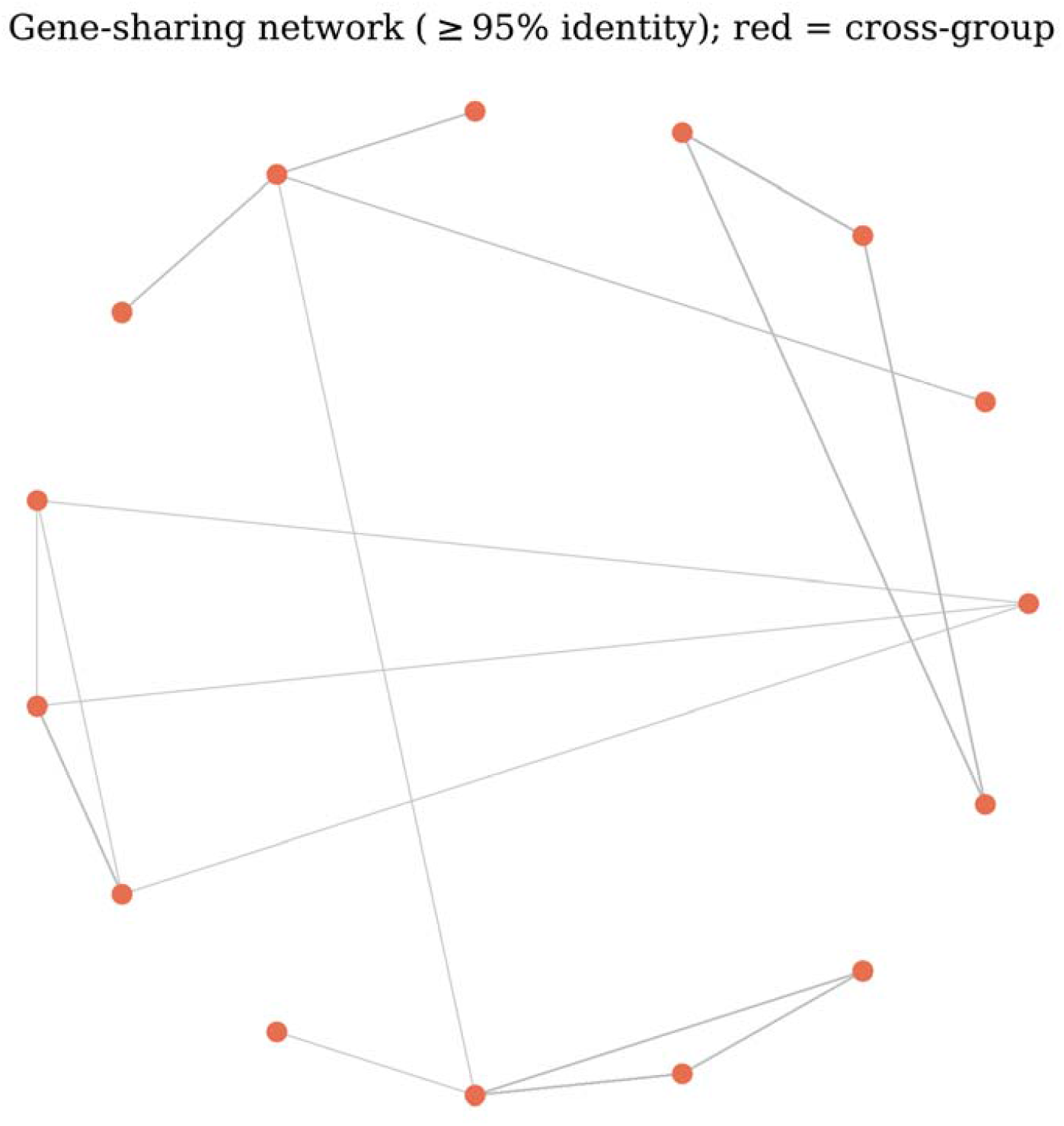
Gene-sharing network. Edges connect genomes sharing an ARG at ≥95% nucleotide identity; nodes are coloured by group. All edges fall within the pathogen/comparator group; no cross-group (red) edge is present.

**Supplementary Fig. 6.**
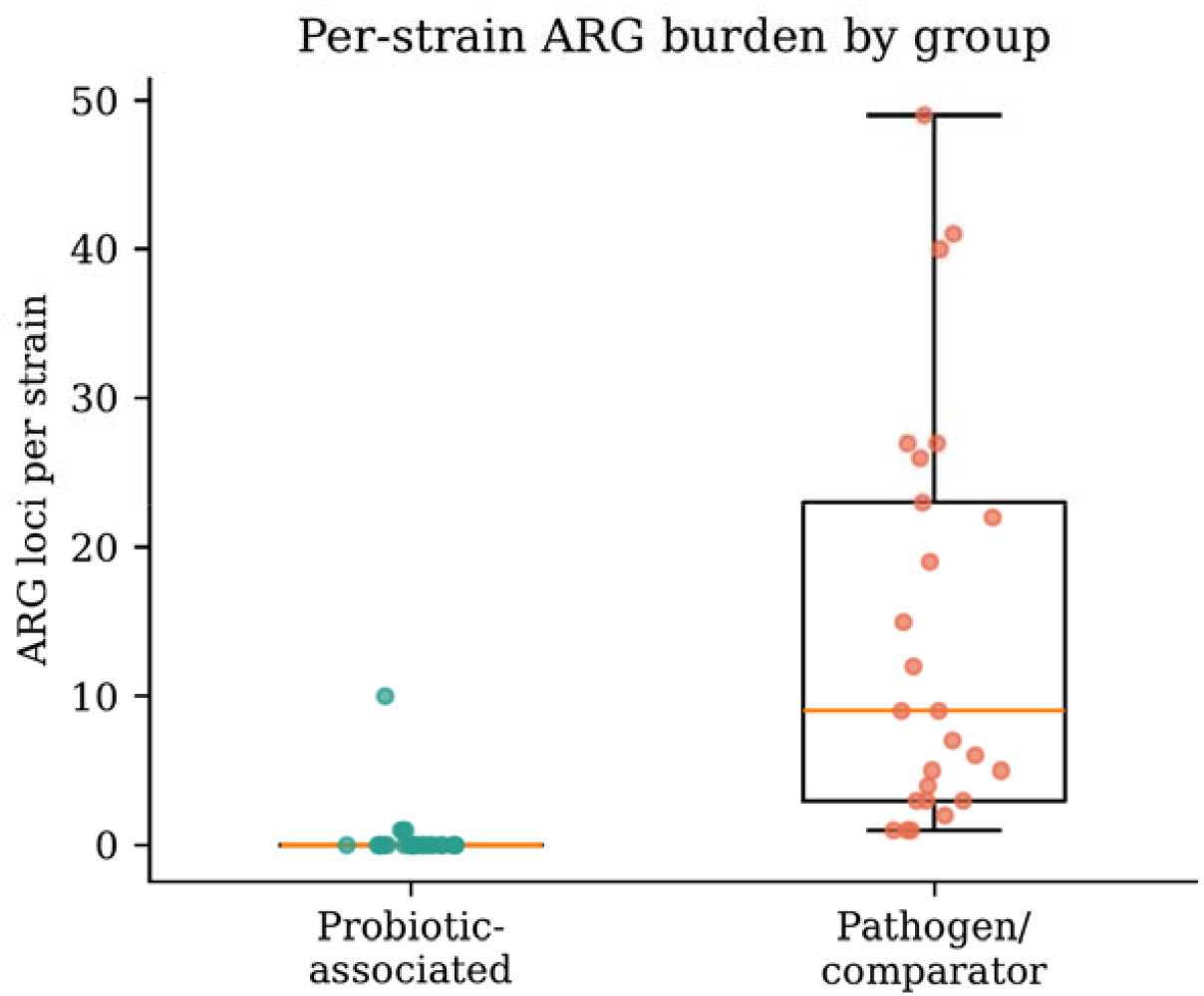
Per-strain ARG burden by group. Boxplots with per-strain points; pathogen/comparator genomes carry substantially more ARG loci than probiotic-associated genomes (Mann–Whitney U = 606.5, P< 0.001).

**Supplementary Fig. 7.**
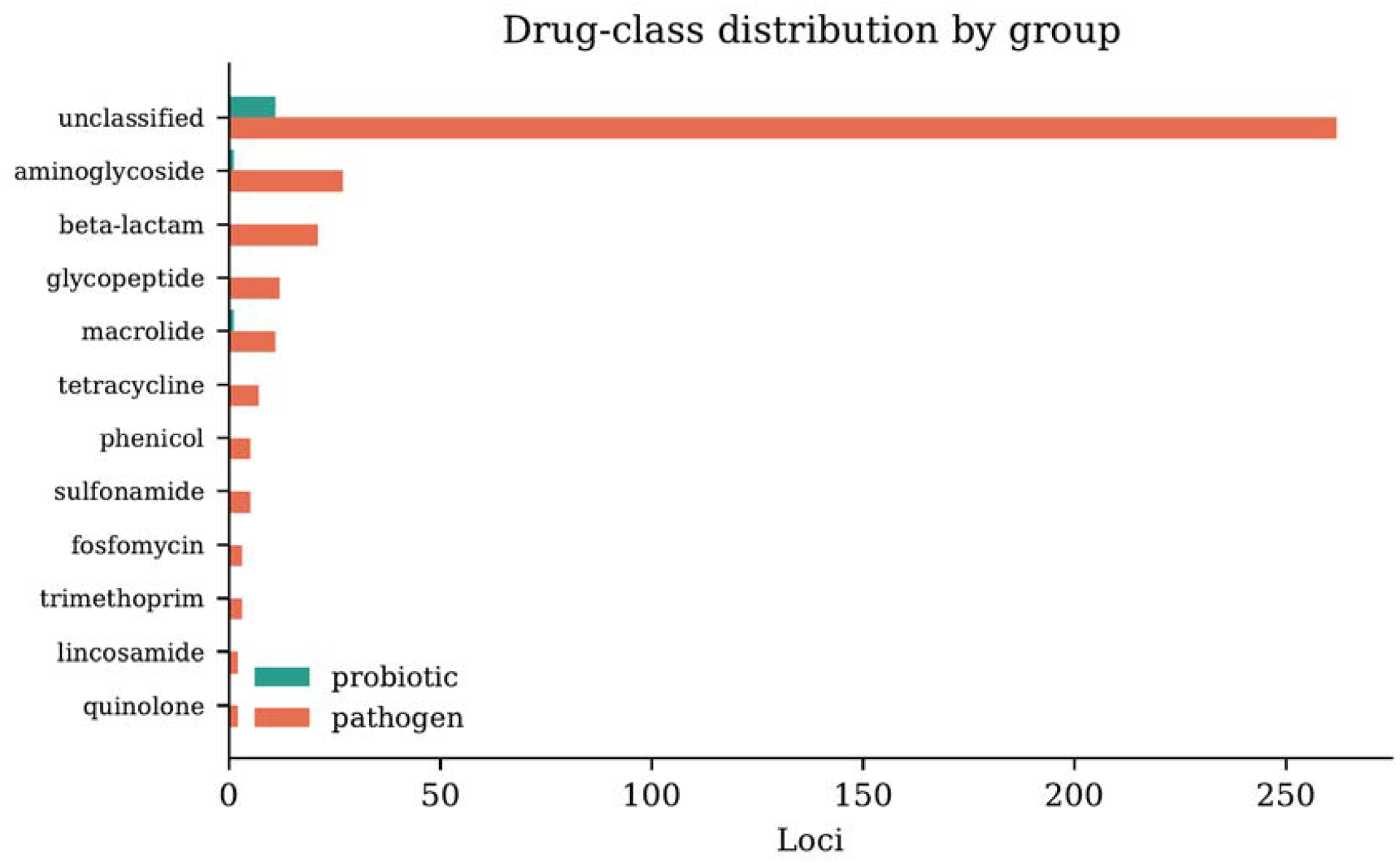
Drug-class distribution by group. ARG loci by drug class, split by group; acquired resistance classes are concentrated in pathogen/comparator genomes.

